# How Not to be Seen: Predicting Unseen Enzyme Functions using Contrastive Learning

**DOI:** 10.64898/2026.02.23.707489

**Authors:** Xiang Ma, Parnal Joshi, Iddo Friedberg, Qi Li

## Abstract

**Motivation:** Predicting enzyme function from its sequence is still an unsolved problem in the life sciences. Moreover, with the explosion of annotated genome data, we are inundated with potential enzymatic sequences that have not yet been biochemically characterized. While it is not possible to assign a not-yet-existing label to such a sequence, there is high value in placing the sequence as accurately as possible in known function space. Doing so can help provide more accurate falsifiable hypotheses for experimentalists wishing to characterize enzymes from specific functional families.

**Results:** Here we present a contrastive learning algorithm for predicting enzyme function from sequence. Our method, EnzPlacer, predicts the third, second, and first EC numbers for a protein whose fourth EC number is not in the training corpus. This novel prediction mechanism accurately places a protein sequence within a narrowed-down functional context, even if the precise function remains unknown.

**Availability:** EnzPlacer is available from https://github.com/drxiangma/EnzPlacer under a GPL3 license.

## Introduction

Protein function prediction is a crucial and yet unresolved problem in computational biology[10, 15]. While experimental annotations curated manually from literature are the most precise way of associating a protein with its function(s), this is time-consuming, and fewer than 0.1% of publicly available protein sequences are annotated that way [28]. We therefore turn to computational protein function prediction to fill in the gaps. Over the years, many different protein function prediction models have been proposed, employing varied techniques such as homology, Hidden Markov Models, text mining, neural networks, and protein language models, to name a few[30, 13, 19, 15, 21].

The representation of protein function is typically done in an ontology [11]. A function ontology, such as the Gene Ontology [3], the Human Phenotype Ontology [23], or the Enzyme Commission Classification (EC) [18], typically describes functional terms that are connected to each other via formally defined relations. This relationship is a directed acyclic graph (DAG), and function prediction models therefore aim to associate one or more sub-DAGs from an ontology with a protein sequence. In this study, we limit our scope to enzymes and the EC ontology. The EC ontology classifies enzymes hierarchically, where the EC class (1^st^ level) identifies the type of reaction (e.g., oxidoreductases); subclass (2^nd^) specifies the type of substrate or group transformed; sub-subclass (3^rd^) details the reaction mechanism, and serial number (4^th^) uniquely identifies the specific substrate/enzyme activity.

Typically, function prediction models aim to solve a structured-output learning problem: associate one or more sub-DAGs of the ontology with a protein sequence [9, 30, 19]. However, high-throughput genomic sequencing is inundating us with putative enzyme sequences that require interpretation. In many cases, those enzymes will have yet unseen functions. That is, we have not yet encountered an enzyme that performs the exact function in nature, and therefore, an EC serial number does not exist for it. In other words, there is no precise label to classify it at the fourth position. However, there is considerable utility in placing the unseen function in the 3^rd^ or even 2^nd^ EC position, which will provide us with a narrower scope of its putative functions [20]. Even if our understanding of the function is imperfect, having an approximate idea rules out certain possibilities and can provide directions for which experiments to run to give a more precise functional characterization.

Here we present a study of how well enzyme function predictors perform when provided with unseen sequences and unseen EC classes. Specifically, we examine how accurately different methods predict the function of an unseen enzyme. The method we developed for this task, EnzPlacer, outperforms existing methods in this task. As genomic data continue to accumulate and require functional annotations, methods that predict as closely as possible to unseen functions will become increasingly necessary to help synergize the utility of computational and experimental methods.

## Methods

### Data

We collected 237,492 enzyme records from the ExPASy ENZYME database [4] (09-Apr-2025 release). We mapped ENZYME entries to UniProtKB accessions to retrieve amino-acid sequences [1]. Then, we applied a deterministic EC cleaning pipeline to retain a single EC assignment per unique protein sequence. First, we removed multifunction enzymes, i.e., proteins annotated with two or more distinct EC numbers, which eliminated 13,247 proteins. We then filtered out proteins with incomplete EC strings (not all four EC fields present) or invalid EC annotations after normalization, including non-digit characters after stripping optional “EC” prefixes/whitespace and any EC containing “99”, removing an additional 7,331 proteins. Finally, we deduplicated EC-sequence pairs by keeping only one record when multiple entries shared the same EC label and identical sequence, removing 33,301 proteins. After cleaning, the dataset contained 183,613 proteins.

#### Unseen Data

Our goal is to simulate a realistic application scenario: newly emerging (previously unseen) EC classes often have only a small number of proteins, whereas well-established EC classes have abundant proteins. To reflect this head–tail imbalance, we constructed an EC-grouped split at the EC4 level, where “EC4” denotes the full four-field EC (e.g., 2.1.3.4). We refer to the corresponding EC3 family as the first three fields (e.g., 2.1.3). We first moved the EC4 group with the most proteins within each EC3 family into the training set, treating it as the established class. We then targeted an 80/20 train/test split by constructing a test-candidate set (which will be further processed later): among the remaining EC4 groups, we sorted them by ascending number of proteins and added the entire EC4 group to the test candidate set until reaching the target test size. This yielded 146,881 training proteins (567 unique ECs) and 36,732 test-candidate proteins (4,260 unique ECs). By construction, (i) EC labels in the test-candidate set do not appear in the training set, and (ii) every EC3 family appearing in the test-candidate set also appears in training, ensuring that test proteins come from known EC3 families while evaluating generalization to unseen EC classes within those families. Figure 1A illustrates this unseen split.

**Fig. 1.**
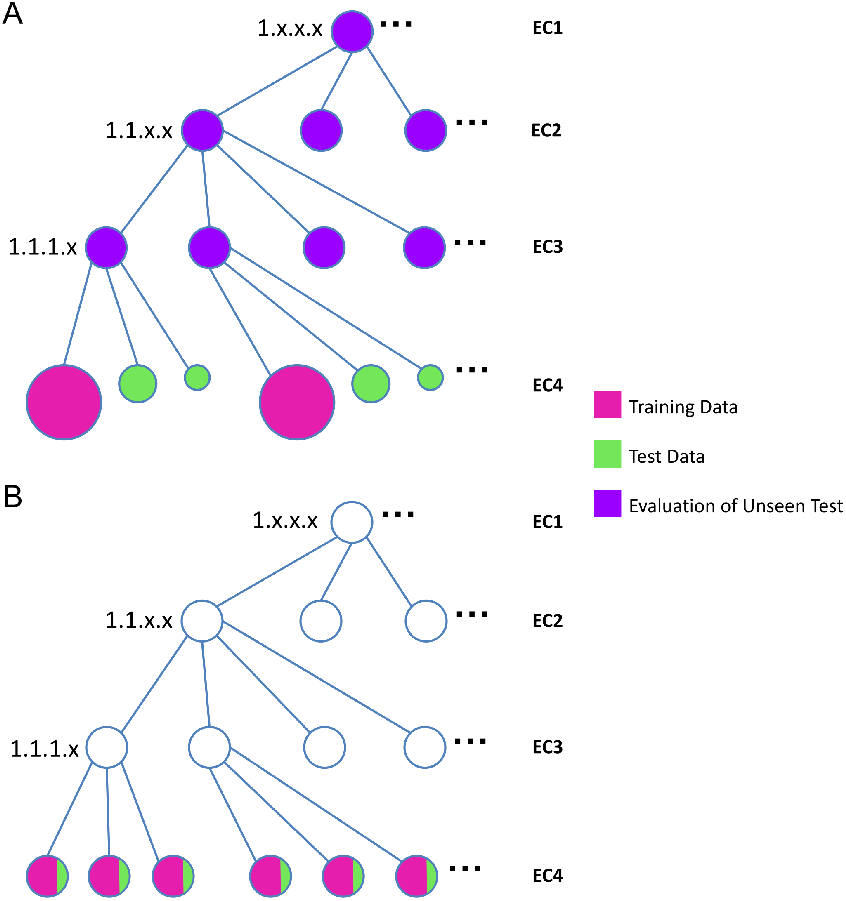
Unseen vs. seen splitting strategies over the EC hierarchy. **(A)Unseen split:** EC labels in the test set do not appear in training, while all EC3 families in test exist in training. We evaluate on the EC1–3 families. Note that the leaf node size reflects the head–tail imbalance: the EC classes containing more proteins within an EC3 are assigned to training (larger leaves), whereas the EC classes containing fewer proteins are in test (smaller leaves). **(B) Seen split (traditional):** All EC labels in the test set exist in training (in-distribution evaluation).

To better reflect real-world evaluation, we further refined the test-candidate set into an experimentally validated subset. In practice, many protein–EC annotations are inferred computationally or propagated by homology, so evaluating on the full test-candidate set can conflate true functional generalization with label noise or uncertain assignments. For our task, ground-truth reliability is especially important: if an EC label is weakly supported, a model may be unfairly penalized (or rewarded) for predicting an alternative but plausible function. We therefore identified experimentally supported EC annotations using evidence metadata (ECO-style evidence in UniProt records) and performed an inner join with the test-candidate set on (*Entry, EC number, Sequence*), yielding an experimentally validated `Unseen Test` set with 9,901 proteins (3,682 unique ECs). Note that, under this split strategy, the training set contains substantially more sequences than the test set, but it covers a much smaller set of unique EC labels than the test set.

To reduce the impact of close homologs, we additionally formed three more challenging low-similarity subsets by filtering protein sequences in the experimentally validated Unseen Test set against the training protein sequences using BLASTp [2]. For each protein sequence, we removed it if any training hit satisfied both (a) percent identity ≥ 50%, 30%, 10% and (b) alignment coverage ≥ 80% on either the query or the subject. This produced three subsets: `Unseen Test 50%` (9,617 proteins; 3,390 unique ECs), `Unseen Test 30%` (8,055 proteins; 3,056 unique ECs), and `Unseen Test 10%` (5,059 proteins; 2,028 unique ECs).

#### Seen Data

To complement the above out-of-distribution evaluations (unseen EC4 within known EC3 families), we also wanted to measure our model’s performance under the traditional in-distribution setting used by many prior studies, where the EC labels in the test set already exist in training. Accordingly, we constructed a seen dataset from the cleaned dataset: we randomly sampled 20% of sequences as candidate test examples and used the remaining 80% as the training pool, which yielded a training set with 146,893 proteins (4,500 unique EC labels). We further refined the candidate test set in two steps: (i) we retained only sequences whose EC labels also appear in the training pool, ensuring a true seen-label evaluation; and (ii) to reduce sequence identity, we applied the same BLAST-based filtering at the 50%, 30%, and 10% identity thresholds, yielding `Seen Test 50%` with 3,022 proteins (1,063 unique EC labels), `Seen Test 30%` with 476 proteins (220 unique EC labels), and `Seen Test 10%` with 212 proteins (102 unique EC labels). Figure 1B illustrates this seen split.

### Model and Protein Representation

Given a protein sequence *x*, we obtain a pretrained embedding *h*(*x*) ∈ ℝ^*d*^ from ESM-1b [22]. We train a lightweight projection head *f*_*θ*_ (a multi-layer perceptron, MLP) to produce the representation used for classification:

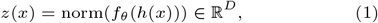

where norm(·) denotes ℓ_2_ normalization and *d* and *D* are the embedding and projection dimensions, respectively.

### Hierarchical Exemplar Contrastive Objective (HiNCE)

Contrastive learning trains an embedding model by *comparing* examples: it pulls representations of similar items closer together and pushes representations of dissimilar items farther apart. In supervised settings, “similar” typically means sharing the same label. This approach has recently shown strong performance for enzyme function transfer by learning a representation space where proteins with the same EC annotation cluster, while proteins from different functions separate. For example, CLEAN [29] applies supervised contrastive learning to improve neighborhood quality beyond raw pretrained embeddings. However, standard supervised contrastive objectives treat the EC label as a *flat* class and do not explicitly enforce the hierarchical structure of the EC system. In our setting, the key goal is not only to separate different EC classes, but also to preserve *hierarchy-consistent neighborhoods* so that proteins remain close when they share the same EC3/EC2/EC1 prefix even if their EC4 differs. Thus, we propose HiNCE, which augments the previous supervised contrastive learning by a hierarchical exemplar term inspired by exemplar-wise contrastive learning [16].

We start with introducing the supervised contrastive loss. During training, each protein serves as a reference (anchor). For each anchor embedding *z*_*a*_, we sample a set of positives 𝒫_*a*_ (proteins sharing the same EC label as the anchor) and a set of negatives 𝒩_*a*_ (proteins from different EC families), and denote *S*_*a*_ = 𝒫_*a*_ ∪ 𝒩_*a*_ (the set of sampled positives and negatives for anchor *a*). The instance-level supervised contrastive loss is:

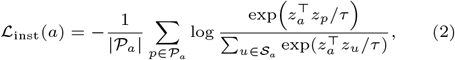

where | · | denotes the number of elements, and *τ* is the temperature parameter (set to 0.1 in all experiments, following CLEAN [29]). This term encourages each anchor to be close to its sampled positives while being separated from sampled negatives, improving local neighborhood discrimination.

To explicitly incorporate EC hierarchy, HiNCE adds an exemplar term that pulls protein embeddings toward *hierarchy-consistent centroids*. For each prefix depth ℓ ∈ *{*1, 2, 3, 4*}* and each prefix class *c*, we compute a centroid as the mean of embeddings belonging to that class:

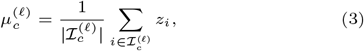

where 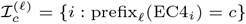. For example, if ℓ = 3 and *c* = `2.1.3`, then 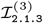 is the set of embedding indices whose EC labels begin with `2.1.3`.

To avoid trivial self-influence especially when a class has few members, for the class with anchor *a*, we use a leave-one-out (LOO) centroid for that class:

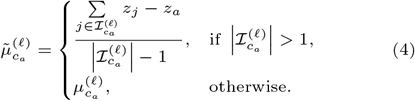

The exemplar-wise NCE loss at each hierarchy depth is then defined as a softmax classification against all centroids at that depth:

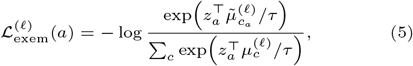

where the denominator sums over all ECℓ prefix classes *c*. Averaging across depths yields

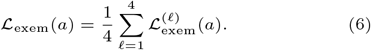

Intuitively, for an enzyme labeled `1.2.3.4`, this exemplar term encourages its embedding to align with the centroids of `1` (EC1), `1.2` (EC2), `1.2.3` (EC3), and `1.2.3.4` (EC4), while separating it from centroids under different EC classes such as `2.*.*.*`. This explicitly shapes the global geometry so that neighborhoods remain informative at higher EC levels even when EC labels differ, which is crucial for our unseen-EC4 evaluation.

Finally, HiNCE combines the local instance discrimination and hierarchical exemplar structuring:

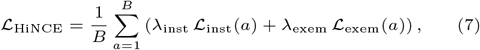

where *B* is the number of embeddings in the iteration, and *λ*_inst_ and *λ*_exem_ control the trade-off between local contrastive separation and hierarchy-aware global organization.

### Training and Testing

#### Hard Negative Samples

Following CLEAN [29], we emphasize hard negatives, which are proteins that do not share the anchor’s EC label but are currently most similar to the anchor in the learned embedding space. Concretely, we periodically compute embeddings for all training proteins under the current model and build a distance map that stores, for each training protein, a shortlist of its nearest neighbors ranked by embedding similarity (equivalently, smallest distance since embeddings are ℓ_2_-normalized). During subsequent iterations, negative samples are preferentially drawn from this shortlist, ensuring the model focuses on separating confusable enzyme families rather than only easy negatives.

Because the embedding space evolves during training, a fixed distance map quickly becomes stale. We therefore refresh the distance map at a fixed interval by recomputing training embeddings and updating each protein’s nearest-neighbor shortlist.

#### Implementation

As input to all models, we computed fixed-length protein embeddings using a pretrained ESM encoder [22]. Unless otherwise stated, we treat these embeddings as frozen features and do not update the pretrained encoder. For EC classes represented by only one sequence in the training set (12 such proteins; 0.0082% of training proteins), following CLEAN [29], we construct an additional positive view by mutating the sequence: we replace a randomly chosen residue with a special mask character (“`*`”) and recompute its pretrained ESM embedding while keeping the encoder fixed. The embedding of the original sequence and the embedding of its masked variant are then treated as a positive pair during contrastive training, enabling single-sequence EC classes to contribute a non-trivial positive signal.

We train the embedding projector with mini-batch optimization using Adam. Each training iteration treats every protein embedding in the mini-batch as an anchor, and contrastive pairs are formed by randomly sampling multiple positives from the same EC label and sampling negatives from different EC labels. Both positives and negatives are re-sampled at each iteration, but negatives are drawn from a mined pool of hard negative EC classes, which is refreshed at a fixed interval after recomputing training embeddings and updating the distance map. In addition, HiNCE uses exemplar-based terms whose class means are recomputed from the current mini-batch embeddings, so the exemplar centroid evolves throughout training as the projector parameters update. Hyperparameters (e.g., learning rate, refresh frequency, and the loss weights *λ*_inst_ and *λ*_exem_) are selected on a held-out validation set.

#### Test

At test time, we embed each query protein and assign the query the EC label of its closest training neighbor in the learned space (this can be extended to *k*-nearest-neighbor voting easily). To evaluate, we report performance at EC1–3 by comparing the corresponding EC prefixes of predicted and ground-truth EC labels. For each of EC1–3, we compute accuracy and macro-averaged precision, recall, and F1 (the harmonic mean of precision and recall). We quantify uncertainty using nonparametric bootstrap over test proteins (1,000 resamples) and report the bootstrap mean together with a 95% confidence interval (shown as mean ± half-width).

### Baselines

We compare EnzPlacer against representative learning-based enzyme annotation methods (CLEAN [29], GloEC [12], and ProteInfer [26]) and a homology-transfer baseline (BLASTp [2]).

- CLEAN [29] utilizes a contrastive-learning approach, and we choose the SupCon-Hard loss as a strong baseline. In our implementation, we additionally apply a simple post-processing step and keep only the first predicted EC per protein, consistent with our single-label setting after removing multi-function proteins.
- GloEC [12] leverages a hierarchy-GCN (Graph Convolutional Network) encoder to globally model dependencies among enzyme labels on the EC hierarchy graph.
- ProteInfer [26] employs deep dilated convolutional networks to learn a direct mapping from full-length protein sequences to functional annotations, capturing long-range sequence patterns without requiring explicit homology search.
- As a non-learning reference, we use BLASTp [2] homology transfer by assigning each query the EC label of its top hit from a BLAST database built on the training sequences.

Finally, we note that several widely used EC predictors (e.g., ECPred [7], DeepEC [25], and DeepECTransformer [14]) do not release training code that enables retraining on custom splits, making them difficult to evaluate fairly under our dataset construction and unseen-EC4 protocol.

## Results and Discussion

### Unseen Prediction Task Setup

We evaluate the ability of enzyme annotation methods to *place proteins with unseen EC labels* into the correct *known functional neighborhood*. Concretely, train and test are disjoint at EC4, while each EC3 family appearing in the test set is represented in training (Methods). Thus, the key question is whether a model can still reliably predict higher-level EC prefixes (EC1–3) for proteins whose exact EC4 label was never observed. EnzPlacer addresses this by learning a function-aware embedding space with the hierarchical contrastive objective (HiNCE) and performing the nearest-neighbor label transfer in that space (Fig. 2).

**Fig. 2.**
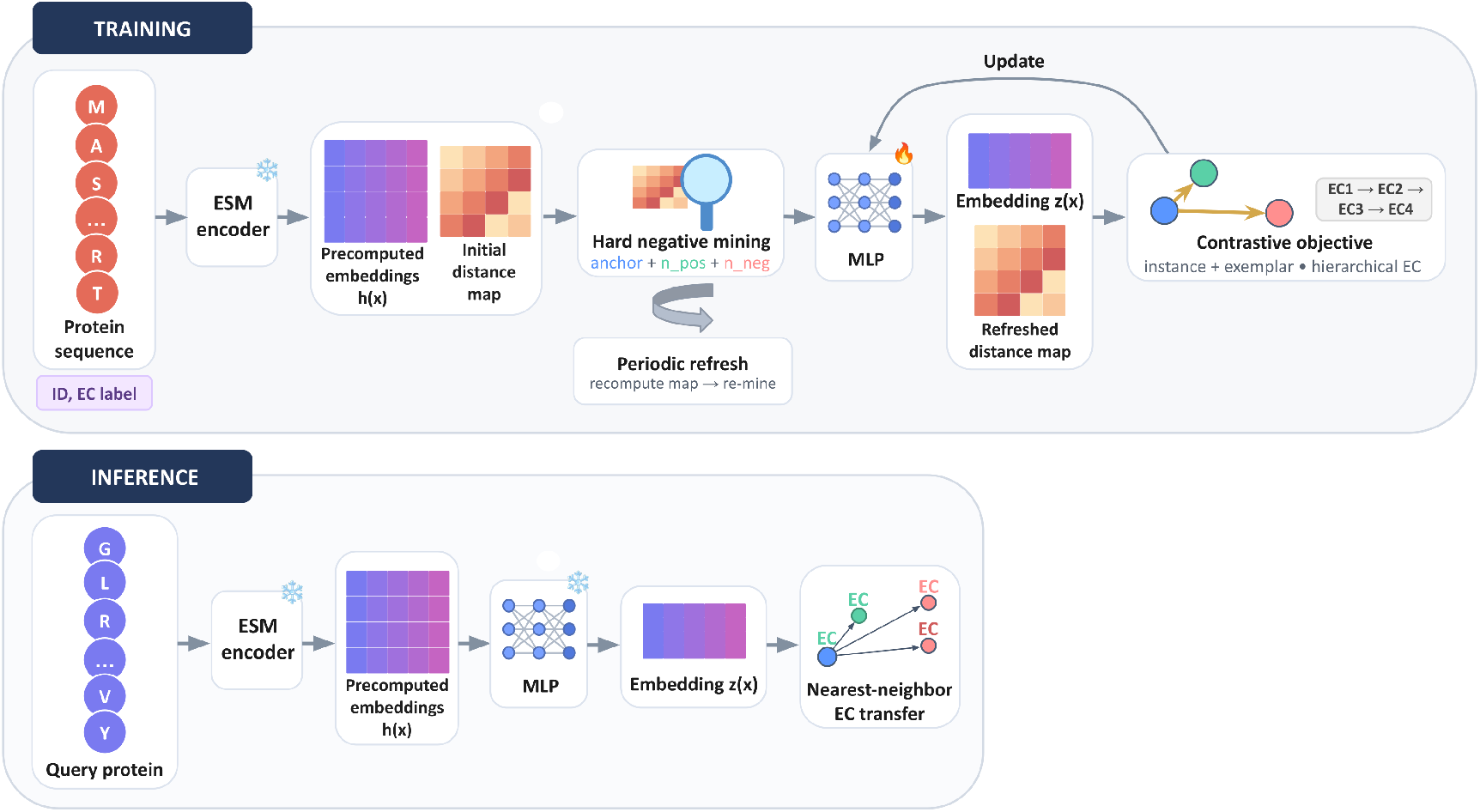
Overview of EnzPlacer. During training, we map fixed ESM embeddings through a projection head (MLP) and optimize the proposed hierarchical exemplar contrastive objective with periodic hard-negative refresh. During inference, a query is embedded and annotated by the nearest reference label in the learned space.

### Unseen Dataset Statistics

We performed a statistical analysis of our dataset to characterize the label-frequency distribution induced by our split construction. Fig. 3A–B show the distribution of the number of sequences per EC in the training and test splits, respectively. In training (Fig. 3A), a large number of ECs have hundreds to thousands of sequences, while the test split (Fig. 3B) is more concentrated at the extreme low-count regime, with most ECs represented by only a handful of sequences. This is expected from our EC4-grouped splitting strategy, which assigns rarer EC4 groups to the test set. Consequently, test examples are enriched for low-frequency, harder-to-generalize ECs, which directly aligns with our objective of evaluating unseen-EC4 generalization. Fig. 3C summarizes the distribution of sequences across EC level-1 classes for both splits. The overall composition across the seven top-level EC classes is broadly consistent between training and test, and certain classes contribute substantially more sequences than others. Together, these observations indicate that our evaluation setting reflects the realistic long-tailed nature of enzyme annotation while deliberately emphasizing difficult, rare EC4 classes in the test set. This makes the unseen-EC4 task a meaningful stress test for whether methods can infer correct higher-level functional context despite sparse labels and limited homology signals.

**Fig. 3.**
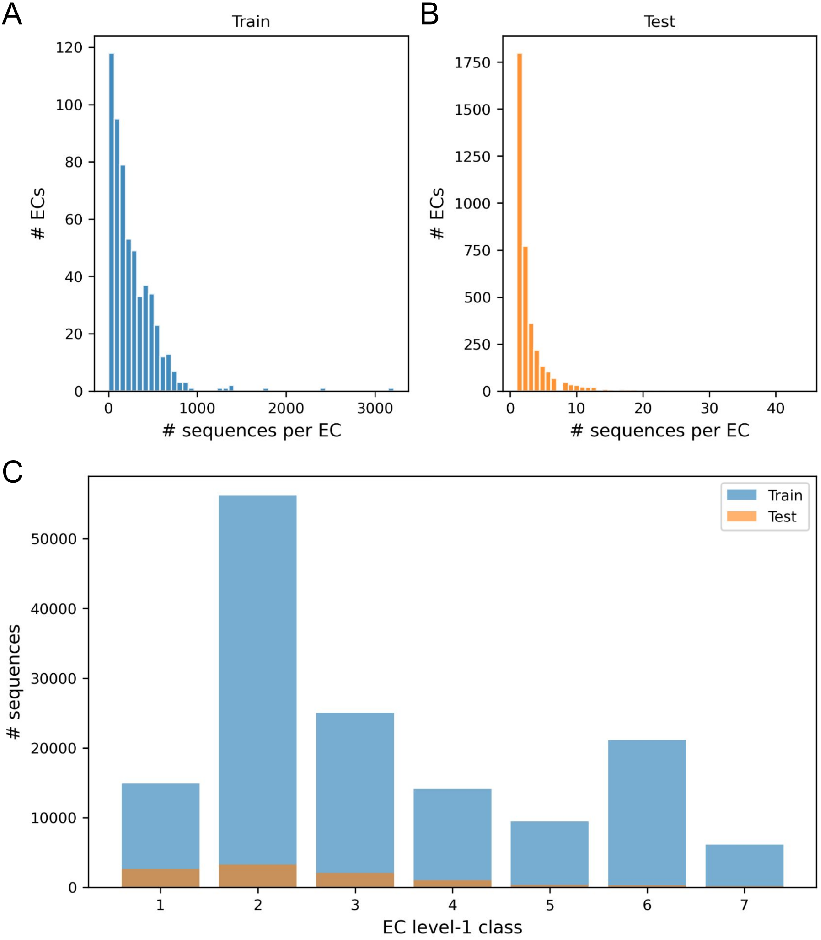
Unseen Dataset statistics. (A–B) Distributions of the number of sequences per EC in the training and test splits highlight the long-tailed label frequency. (C) Sequence counts per EC level-1 class for train and test.

### Unseen-EC4 Result

On the experimentally validated Unseen Test set, we observe that all methods achieve relatively modest scores, highlighting that predicting enzyme function when the exact EC label in the test set is absent from the training label space is intrinsically difficult (Fig. 4A–B). In Fig. 4A (EC2 evaluation), EnzPlacer attains the highest accuracy (0.4350 ± 0.0099) and macro-F1 (0.2614 ± 0.0157), with CLEAN (0.3854 ± 0.0099 accuracy, 0.2261 ± 0.0137 macro-F1) and GloEC trailing behind, while ProteInfer is substantially lower. The same pattern holds in Fig. 4B (EC3 evaluation), where EnzPlacer again leads with 0.3563 ± 0.0096 accuracy and 0.1678 ± 0.0098 macro-F1. Notably, precision and recall follow similar rankings across models in both panels, suggesting that EnzPlacer reduces both false positives and false negatives relative to the baselines, rather than improving performance only through a precision– recall trade-off.

**Fig. 4.**
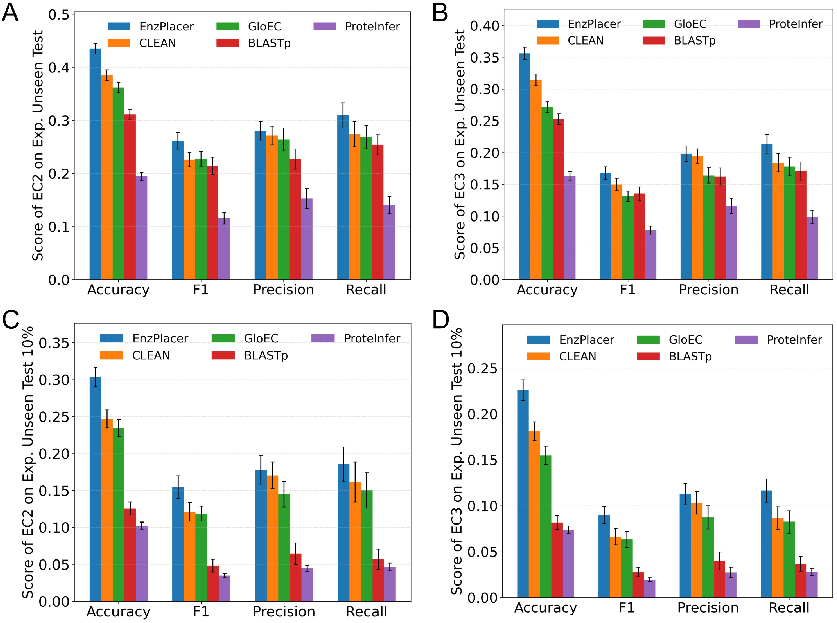
Unseen setting: experimentally validated test results. Performance at EC2 and EC3 on the experimentally validated Unseen Test set, and on the more challenging low-similarity subset (Unseen Test 10%). Error bars indicate mean ± half-width of the 95% bootstrap confidence interval.

From these observations, several key conclusions emerge. First, this is a highly challenging and comparatively under-studied regime: even the best method remains far from saturated performance, especially at EC3 where macro-F1 values are low across the board, suggesting substantial room for methodological advances. Second, EnzPlacer is consistently the strongest across metrics at both EC2 and EC3, indicating that its learned geometry better preserves function-relevant neighborhoods when EC4 labels shift. Third, BLASTp remains a competitive baseline in this setting and performs reasonably well given its simplicity, which is expected because many test proteins may still have detectable homologs in the training database and homology transfer can often recover higher-level EC prefixes. Lastly, while learning-based baselines such as CLEAN and GloEC improve accuracy over BLASTp, their gains in macro-F1 are comparatively smaller, suggesting that class imbalance and difficult minority families remain a major bottleneck in unseen-EC4 generalization.

Fig. 4C–D reports results on a more stringent subset derived from the experimentally validated Unseen Test set by filtering out any test sequence that has a close training hit, leaving only low-similarity cases (here, < 10% identity under our BLASTp criterion). As expected, performance decreases for every method relative to Fig. 4A–B. In Fig. 4C (EC2), EnzPlacer drops from 0.4350 to 0.3034 ± 0.0130 accuracy and from 0.2614 to 0.1547 ± 0.0153 macro-F1, and similar declines are observed for CLEAN and GloEC. In Fig. 4D (EC3), EnzPlacer reaches 0.2260 ± 0.0112 accuracy and 0.0899 ± 0.0091 macro-F1. A striking pattern in Fig. 4C–D is that BLASTp degrades sharply compared to the other approaches, consistent with the intuition that homology transfer is fragile when near-neighbor sequence similarity is explicitly removed. Similar patterns are observed for the low-similarity subsets obtained using the *<* 50% and *<* 30% identity thresholds. Detailed results across all metrics at EC1–3 for `Unseen Test, Unseen Test 50%, Unseen Test 30%,` and `Unseen Test 10%` are provided in Supplementary Table S1.

The sharp BLASTp collapse under low similarity supports the claim that the remaining test proteins require signals beyond nearest-homolog transfer. In contrast, although EnzPlacer and the learning-based baselines also decline, their relative drops are smaller, indicating that learned representation spaces can retain some function-relevant structure even when sequence identity is extremely low. Importantly, EnzPlacer preserves the strongest performance margin over baselines at both EC2 and EC3 in this harder regime. This suggests that the hierarchy-aware training signal helps sustain informative neighborhoods when the most direct homology cues are absent.

We identified multiple phosphodiesterases for which EnzPlacer correctly recovers the EC3-level functional annotation, while baseline methods including BLASTp and embedding-based approaches without hierarchical objectives incorrectly place the proteins into unrelated functional classes. For example, protein `A0A1D8PNZ7` has the experimentally verified annotation EC `3.1.4.2` [5]. However, this sequence, when BLASTed (BLASTp) against the training set, is assigned to EC `2.7.11.1`, protein kinase, reflecting a failure to predict even coarse-grained functional chemistry with low sequence similarity. Similarly, other embedding baselines such as CLEAN and GloEC predict EC subclasses associated with transferases, which do not share phosphodiester bond hydrolysis as their defining reaction. In contrast, EnzPlacer places this protein within EC `3.1.4`, predicting `3.1.4.11`, thereby preserving the correct EC3 neighborhood despite the EC4 label being unseen during training. A similar pattern is observed for `B8NQ51` (true EC `3.1.4.41`) [8], where BLASTp and baseline embedding methods again drift to unrelated EC2 or EC3 families, while EnzPlacer correctly assigns a phosphodiesterase EC3 label. In these cases, EnzPlacer’s nearest neighbors in the learned embedding space are enriched for enzymes that share phosphodiester bond hydrolysis activity, even when sequence identity is low and EC4 serial numbers differ.

Taken together, Fig. 4 demonstrates that the experimentally validated unseen-EC4 evaluation is a demanding benchmark for enzyme function placement: all models struggle, and performance deteriorates further when close homologs are removed, underscoring the difficulty of generalizing to novel EC4 serial numbers and distant sequences. Within this challenging landscape, EnzPlacer achieves the best results across all reported metrics at both EC2 and EC3, consistent with its objective of preserving *hierarchy-consistent neighborhood structure*. Specifically, the instance-level supervised contrastive component enforces local discrimination among nearby proteins, while the hierarchical exemplar term promotes global organization across EC levels. In practice, this combination helps identify neighbors that remain consistent at EC3/EC2 even when EC4 labels differ, aligning the learned geometry with the EC ontology and improving higher-level functional placement. BLASTp remains a strong but similarity-dependent baseline, but its limitations become clear under stringent similarity filtering. Finally, the modest macro-F1 scores across methods, especially at EC3, highlight that unseen-EC4 function placement remains far from solved, motivating future work on richer hierarchy-aware objectives, improved handling of rare EC families, and stronger integration of complementary functional signals beyond sequence identity alone.

### Seen Prediction Task Setup

In the seen setting, our goal is to accurately predict EC labels when the test label space is contained within the training label space. Similarly to the unseen setting, we learn a projection network (MLP) that maps fixed protein embeddings from ESM into a task-specific representation space, and we optimize this space with contrastive learning such that proteins sharing the same EC label are pulled together while proteins with different EC labels are pushed apart. Our EnzPlacer additionally encourages hierarchical organization by further clustering EC labels that belong to the same EC1–3 families, promoting family-consistent structure in the embedding space. At inference time, we transfer EC labels from the nearest neighbor in this learned space and evaluate the predicted EC labels at EC4 level.

### Seen Dataset Statistics

We analyzed the label-frequency and class-composition statistics of the seen dataset (Seen Test 50%), with detailed results summarized in Supplementary Figure S1. Compared with the unseen split, the distribution of sequences per EC label is more similar between the training and test splits, consistent with the fact that the seen split is not explicitly constructed to concentrate the test set in the low-count regime. At the same time, the distribution of sequences across EC level-1 classes remains imbalanced in both training and test, mirroring the pattern observed in the unseen setting where certain EC1 classes contribute substantially more sequences than others.

### Seen-EC4 Result

To evaluate the models under the traditional in-distribution task used by many prior studies, we report results in a seen setting (Seen Test 50%), where EC labels in test are present in the training label space (Fig. 5). In this setting, EnzPlacer achieves an accuracy of 0.9098 ± 0.0074 and a macro-F1 of 0.7529 ± 0.0144, exceeding BLASTp (0.8971 ± 0.0071 accuracy; 0.7342 ± 0.0138 macro-F1) and CLEAN (0.8775 ± 0.0084 accuracy; 0.7262±0.0141 macro-F1). Precision and recall follow the same ordering, and hierarchical accuracies are uniformly high, with Acc EC1–EC3 all above ∼0.96 for EnzPlacer (Fig. 5). Similar performance trends and relative rankings are observed for the more stringent `Seen Test 30% and Seen Test 10%` subsets. Detailed results of metrics for `Seen Test 50%, Seen Test 30%`, and `Seen Test 10%` are reported in Supplementary Table S2.

**Fig. 5.**
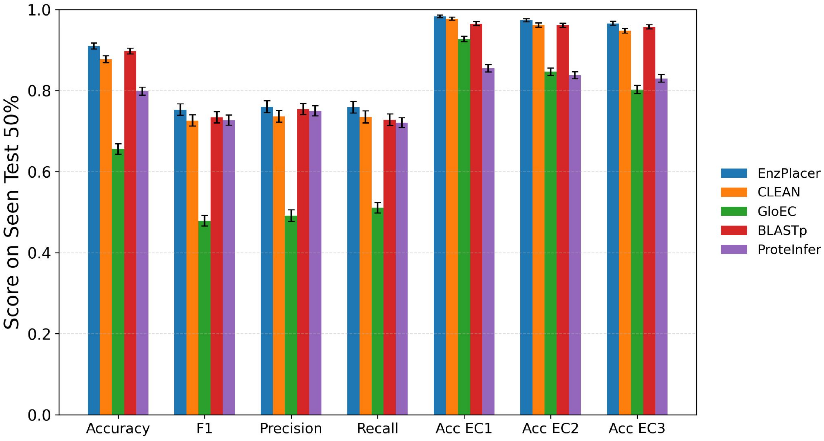
Seen setting: test performance when EC4 labels are present. Results on a “seen” test set (Seen Test 50%) where EC4 labels are drawn from the training label space. Bars show mean ± half-width of the 95% bootstrap confidence interval.

Overall, these results align with the expected behavior of the traditional seen-label regime. First, all methods perform substantially better than in the unseen-EC4 evaluation, with markedly higher accuracy and macro-F1 across the board, consistent with prior reports that learned-embedding retrieval and homology transfer are highly effective when test labels are already represented during training. Second, BLASTp remains a particularly strong baseline, underscoring the continued value of homology-based transfer under in-distribution assumptions. Third, accuracy at EC1–3 levels is near-saturated (Fig. 5), indicating that coarse-grained functional assignment is highly reliable when EC labels are present and that EnzPlacer’s gains are achieved without sacrificing higher-level correctness.

Importantly, the sharp contrast between Fig. 5 and the unseen-EC4 results in Fig. 4 highlights the core challenge studied in this work. When EC labels are present, multiple methods perform well and approach saturation. However, when EC labels are held out, performance drops substantially for all models, especially at EC3. This gap underscores that unseen-EC4 enzyme function placement remains a demanding and comparatively under-explored benchmark, and it motivates broader community attention toward hierarchy-aware objectives and evaluation protocols that explicitly target generalization beyond observed EC4 serial numbers.

### Embedding Visualization

To visualize how representation learning changes the geometry of the embedding space, we projected high-dimensional protein embeddings from the Unseen Test set into 2D using t-distributed stochastic neighbor embedding (t-SNE) and colored points by representative EC3 families (Fig. 6). To make a 2D projection, given an embedding vector for each protein, t-SNE constructs a 2D layout by matching local neighborhood relationships: pairs of proteins that are close in the original embedding space are encouraged to remain close after projection, making the visualization especially informative for comparing local clustering structure between representations [17]. We selected eight EC3 families as representative examples. Note that each EC3 family can contain many distinct EC4 groups. For instance, the 3.1.4 EC3 family contains 23 unique EC4 groups in our test set, and under our split strategy many of these EC4 groups have only a small number of proteins. EC 3.1.4 is the Phosphoric diester hydrolase or Phosphodiesterase family, which includes enzymes that catalyze hydrolysis of phosphodiester bonds. While enzymes from this class share core biochemistry, they are heterogeneous in sequence and substrate specificity. As a result, EC 3.1.4 represents a particularly challenging family for annotation transfer, especially when exact EC4 labels are unseen and close homologs are absent. This intra-family heterogeneity is reflected in the raw ESM visualization (top), where the green 3.1.4 points appear widely scattered and inter-mixed with other families. In the EnzPlacer visualization (bottom), the same green family becomes noticeably more concentrated, and a similar trend holds for most other colored EC3 families, indicating improved family-level neighborhood coherence.

**Fig. 6.**
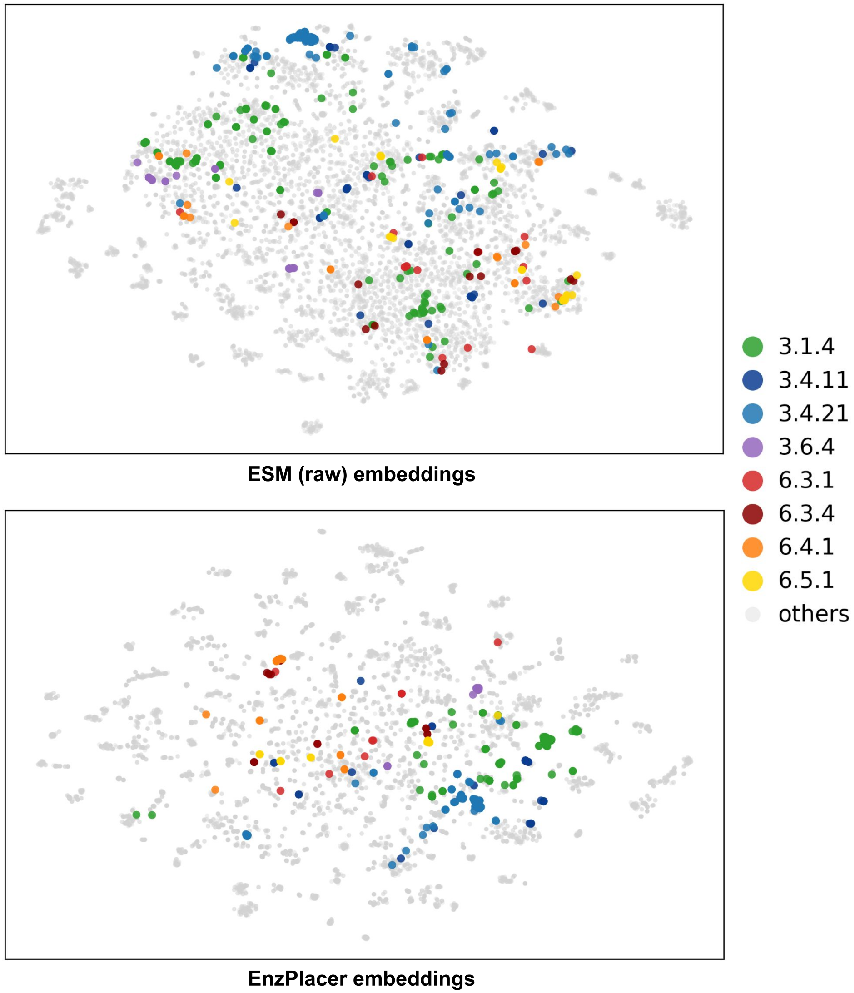
Embedding visualization. Two embeddings were visualized on Unseen Test, including the raw ESM embeddings (top) and the EnzPlacer embeddings (bottom). For readability, we highlight eight representative EC3 families in color and plot the remaining families in gray. EnzPlacer provides a better separation and in-group clustering of the 6.-.- and the 3.-.-enzyme classes than the raw ESM embeddings.

At higher hierarchy levels (EC1–2), the labels are inherently broader because they pool functionally diverse subfamilies across many EC3/EC4 groups. As a result, clustering is expected to be less defined, and different EC1/EC2 families may span wide and partially overlapping regions in 2D. Nevertheless, several hierarchy-consistent patterns emerge when comparing the two panels. At the EC2 level, the sibling families `3.4.11` and `3.4.21` (both within `3.4.*`) appear closer and more mutually coherent in the EnzPlacer space than in raw ESM, consistent with encouraging family-aware organization. Meanwhile, `6.3.1` and `6.3.4` (within `6.3.*`) remain comparatively dispersed, suggesting that some EC2 families remain challenging to consolidate. At the EC1 level, the `3.*.*` groups (cool colors) and the `6.*.*` groups (warm colors) appear more co-located within their respective superclasses and better separated from each other in EnzPlacer than in raw ESM, indicating clearer coarse-level structure.

Overall, these visualizations suggest that EnzPlacer reshapes the embedding space to better align with the EC hierarchy, supporting the interpretation that HiNCE encourages hierarchy-aware neighborhood geometry. In particular, EnzPlacer improves EC3 family coherence even when an EC3 family contains many unseen EC4 labels (e.g., `3.1.4`), which is favorable for neighborhood-based inference. At the same time, the Unseen Test setting remains challenging: the model has not seen either the test protein sequences or the EC labels. As a result, clusters are still partially mixed and not cleanly separated at every level (especially at EC2), leaving room for future improvements in learning sharper hierarchical structure under unseen-label generalization.

### Implications, Limitations and Future Directions

Accurate placement of enzymes at the EC3 level is biologically meaningful because the first three positions of the EC hierarchy denote the core reaction chemistry: class, subclass, and sub-subclass. From a practical standpoint, accurate placement at EC3/EC2 for proteins with unassigned EC4 labels can directly support experimental design to provide a more precise understanding of the enzyme’s function. Rather than attempting to “guess” a novel EC4 serial number, EnzPlacer provides a hierarchy-consistent *functional neighborhood* that narrows plausible reaction mechanisms and substrates. For example, distinguishing a phosphodiesterase from a kinase, esterase, or peptidase constitutes a substantial reduction in functional uncertainty. This is particularly valuable for metagenomic and genome-mining pipelines, where large numbers of putative enzymes lack close homologs and where even a coarse-grained EC3 hypothesis can substantially reduce downstream experimental search space. However, EC3 placement remains challenging, as enzymes that share the same reaction chemistry can arise through convergent evolution, leading to substantial heterogeneity in sequence, domain architecture, and structural context [24, 6]. Consequently, homology-based annotation transfer can fail even at relatively coarse functional levels when sequence identity is low. Additionally, several EC classes encompass large and functionally diverse families in which the defining reaction chemistry is preserved, but substrates, cofactors, or biological roles vary widely. This further weakens the correspondence between sequence similarity and enzyme function class, complicating efforts to reliably group enzymes based on sequence similarity alone [27, 10].

Our study addresses this challenge by focusing on hierarchy-consistent functional placement rather than exact EC4 recovery. Conceptually, our results suggest that the key difficulty in the unseen-EC4 regime is not only representation quality, but *geometry*: the embedding space must preserve hierarchy-consistent neighborhoods so that retrieval remains informative after EC4 labels shift. Prior contrastive-learning approaches for enzyme annotation (e.g., CLEAN) demonstrate that supervised contrastive objectives can improve EC transfer beyond homology [29]. EnzPlacer extends this idea with an explicit hierarchical objective (HiNCE) that injects global structure via level-wise centroids and enforces alignment across EC levels, which better matches our evaluation goal: predicting correct EC prefixes when the exact EC4 serial number is absent. The strong degradation of BLASTp under stringent identity filtering further highlights that, in low-similarity conditions, high-level functional placement requires signals beyond nearest-homolog transfer, and that learning a function-aware metric space can improve robustness (Fig. 4).

There are several practical opportunities for extension. EnzPlacer is designed for *placement* into known functional neighborhoods rather than proposing brand-new EC labels, so its outputs should be interpreted as structured hypotheses that prioritize plausible EC2/EC3 contexts. Our current experiments also focus on single-label enzymes (Methods). In particular, we treat enzymes as single-label entities despite the prevalence of promiscuous or multifunctional enzymes. We also do not explicitly model substrate specificity, cofactors, or cellular context, and rely primarily on sequence-derived embeddings without incorporating structural, kinetic, or regulatory information that can be critical for resolving fine-grained functional differences. Extending the objective and inference to handle multi-label and promiscuous enzymes is an important next step. In addition, while we treat pretrained ESM embeddings as fixed features, integrating complementary signals such as structure-derived representations or curated functional prototypes may further strengthen robustness under extreme sequence divergence. Finally, incorporating calibrated uncertainty and abstention would allow EnzPlacer to better support open-set usage by flagging cases where even coarse-grained placement is unreliable, enabling more effective human-in-the-loop curation and experimental prioritization.

## Acknowledgments

This work is supported in part by funds from the National Science Foundation (NSF: # 1636933 and # 1920920) awarded to QL, by funds from NIH/NIGMS R01GM145937 awarded to IF, and by an Iowa State University Translational AI Center SEED grant awarded to QL and IF.

