## Supplementary Files for "How Not to be Seen: Predicting Unseen Enzyme Functions using Contrastive Learning"


**Table S1. Detailed metrics for unseen evaluations.** Mean ± half-width of the 95% bootstrap confidence interval for EC1–EC3 metrics on `Unseen Test` and the low-similarity subsets (`Unseen Test 50%`, `Unseen Test 30%`, `Unseen Test 10%`).

| Model | Test set | EC1 Acc. | EC1 F1 | EC1 Prec. | EC1 Rec. | EC2 Acc. | EC2 F1 | EC2 Prec. | EC2 Rec. | EC3 Acc. | EC3 F1 | EC3 Prec. | EC3 Rec. |
|---|---|---|---|---|---|---|---|---|---|---|---|---|---|
| BLAST | `Unseen Test` | 0.4894 ± 0.0096 | 0.3563 ± 0.0107 | 0.3586 ± 0.0101 | 0.4075 ± 0.0131 | 0.3112 ± 0.0091 | 0.2143 ± 0.0163 | 0.2270 ± 0.0190 | 0.2541 ± 0.0191 | 0.2524 ± 0.0085 | 0.1355 ± 0.0105 | 0.1621 ± 0.0136 | 0.1708 ± 0.0143 |
| ProteInfer | `Unseen Test` | 0.3983 ± 0.0091 | 0.2956 ± 0.0102 | 0.2879 ± 0.0097 | 0.3461 ± 0.0122 | 0.1943 ± 0.0078 | 0.1156 ± 0.0108 | 0.1529 ± 0.0186 | 0.1405 ± 0.0159 | 0.1627 ± 0.0073 | 0.0779 ± 0.0064 | 0.1160 ± 0.0119 | 0.0986 ± 0.0108 |
| GloEC | `Unseen Test` | 0.6248 ± 0.0096 | 0.5539 ± 0.0140 | 0.5651 ± 0.0148 | 0.5581 ± 0.0155 | 0.3619 ± 0.0098 | 0.2274 ± 0.0137 | 0.2641 ± 0.0215 | 0.2691 ± 0.0215 | 0.2716 ± 0.0086 | 0.1312 ± 0.0082 | 0.1640 ± 0.0120 | 0.1781 ± 0.0146 |
| CLEAN | `Unseen Test` | 0.5950 ± 0.0090 | 0.5257 ± 0.0142 | 0.5497 ± 0.0155 | 0.5323 ± 0.0158 | 0.3854 ± 0.0099 | 0.2261 ± 0.0137 | 0.2714 ± 0.0175 | 0.2739 ± 0.0239 | 0.3144 ± 0.0092 | 0.1498 ± 0.0097 | 0.1945 ± 0.0112 | 0.1841 ± 0.0145 |
| EnzPlacer | `Unseen Test` | 0.6387 ± 0.0093 | 0.5651 ± 0.0128 | 0.5539 ± 0.0141 | 0.5961 ± 0.0143 | 0.4350 ± 0.0099 | 0.2614 ± 0.0157 | 0.2804 ± 0.0177 | 0.3095 ± 0.0237 | 0.3563 ± 0.0096 | 0.1678 ± 0.0098 | 0.1982 ± 0.0123 | 0.2133 ± 0.0153 |
| BLAST | `Unseen Test 50%` | 0.4807 ± 0.0103 | 0.3496 ± 0.0126 | 0.3527 ± 0.0122 | 0.4005 ± 0.0160 | 0.2997 ± 0.0091 | 0.2028 ± 0.0167 | 0.2191 ± 0.0172 | 0.2407 ± 0.0200 | 0.2402 ± 0.0084 | 0.1263 ± 0.0106 | 0.1515 ± 0.0132 | 0.1596 ± 0.0147 |
| ProteInfer | `Unseen Test 50%` | 0.4091 ± 0.0052 | 0.3146 ± 0.0048 | 0.3040 ± 0.0047 | 0.3463 ± 0.0057 | 0.1944 ± 0.0041 | 0.1123 ± 0.0055 | 0.1430 ± 0.0102 | 0.1314 ± 0.0078 | 0.1624 ± 0.0037 | 0.0724 ± 0.0042 | 0.1066 ± 0.0080 | 0.0852 ± 0.0058 |
| GloEC | `Unseen Test 50%` | 0.6189 ± 0.0096 | 0.5467 ± 0.0143 | 0.5590 ± 0.0162 | 0.5512 ± 0.0153 | 0.3522 ± 0.0097 | 0.2130 ± 0.0124 | 0.2507 ± 0.0204 | 0.2578 ± 0.0207 | 0.2601 ± 0.0088 | 0.1212 ± 0.0085 | 0.1548 ± 0.0112 | 0.1548 ± 0.0112 |
| CLEAN | `Unseen Test 50%` | 0.5884 ± 0.0096 | 0.5190 ± 0.0150 | 0.5442 ± 0.0159 | 0.5253 ± 0.0161 | 0.3764 ± 0.0093 | 0.2157 ± 0.0130 | 0.2605 ± 0.0157 | 0.2634 ± 0.0233 | 0.3044 ± 0.0090 | 0.1406 ± 0.0106 | 0.1828 ± 0.0119 | 0.1746 ± 0.0153 |
| EnzPlacer | `Unseen Test 50%` | 0.6329 ± 0.0093 | 0.5587 ± 0.0128 | 0.5475 ± 0.0142 | 0.5905 ± 0.0149 | 0.4267 ± 0.0098 | 0.2512 ± 0.0148 | 0.2694 ± 0.0144 | 0.2999 ± 0.0246 | 0.3469 ± 0.0095 | 0.1597 ± 0.0108 | 0.1873 ± 0.0122 | 0.2052 ± 0.0156 |
| BLAST | `Unseen Test 30%` | 0.4158 ± 0.0114 | 0.2727 ± 0.0266 | 0.2857 ± 0.0266 | 0.3126 ± 0.0328 | 0.2261 ± 0.0093 | 0.1326 ± 0.0163 | 0.1558 ± 0.0211 | 0.1607 ± 0.0226 | 0.1726 ± 0.0081 | 0.0754 ± 0.0093 | 0.0979 ± 0.0134 | 0.0982 ± 0.0130 |
| ProteInfer | `Unseen Test 30%` | 0.3471 ± 0.0057 | 0.2283 ± 0.0056 | 0.2223 ± 0.0050 | 0.2640 ± 0.0089 | 0.1360 ± 0.0041 | 0.0730 ± 0.0064 | 0.1073 ± 0.0112 | 0.0881 ± 0.0096 | 0.1043 ± 0.0036 | 0.0396 ± 0.0027 | 0.0630 ± 0.0067 | 0.0513 ± 0.0058 |
| GloEC | `Unseen Test 30%` | 0.5783 ± 0.0106 | 0.5000 ± 0.0196 | 0.4969 ± 0.0198 | 0.4876 ± 0.0179 | 0.2945 ± 0.0096 | 0.1605 ± 0.0108 | 0.1935 ± 0.0167 | 0.2028 ± 0.0217 | 0.2045 ± 0.0090 | 0.0849 ± 0.0079 | 0.1147 ± 0.0108 | 0.1148 ± 0.0140 |
| CLEAN | `Unseen Test 30%` | 0.5398 ± 0.0113 | 0.4516 ± 0.0161 | 0.4902 ± 0.0167 | 0.4553 ± 0.0205 | 0.3183 ± 0.0099 | 0.1687 ± 0.0114 | 0.2119 ± 0.0146 | 0.2031 ± 0.0212 | 0.2497 ± 0.0093 | 0.0983 ± 0.0075 | 0.1392 ± 0.0111 | 0.1281 ± 0.0134 |
| EnzPlacer | `Unseen Test 30%` | 0.5919 ± 0.0108 | 0.4971 ± 0.0174 | 0.4861 ± 0.0171 | 0.5357 ± 0.0203 | 0.3765 ± 0.0106 | 0.2016 ± 0.0135 | 0.2206 ± 0.0192 | 0.2505 ± 0.0264 | 0.2994 ± 0.0098 | 0.1238 ± 0.0091 | 0.1483 ± 0.0119 | 0.1670 ± 0.0148 |
| BLAST | `Unseen Test 10%` | 0.3239 ± 0.0121 | 0.1821 ± 0.0205 | 0.2033 ± 0.0217 | 0.2042 ± 0.0271 | 0.1255 ± 0.0089 | 0.0483 ± 0.0086 | 0.0644 ± 0.0146 | 0.0572 ± 0.0136 | 0.0818 ± 0.0075 | 0.0280 ± 0.0046 | 0.0398 ± 0.0098 | 0.0367 ± 0.0082 |
| ProteInfer | `Unseen Test 10%` | 0.3251 ± 0.0078 | 0.1804 ± 0.0061 | 0.1826 ± 0.0053 | 0.2044 ± 0.0105 | 0.1023 ± 0.0046 | 0.0348 ± 0.0029 | 0.0446 ± 0.0041 | 0.0465 ± 0.0052 | 0.0738 ± 0.0040 | 0.0195 ± 0.0020 | 0.0273 ± 0.0058 | 0.0280 ± 0.0038 |
| GloEC | `Unseen Test 10%` | 0.5326 ± 0.0142 | 0.3970 ± 0.0260 | 0.4092 ± 0.0267 | 0.4082 ± 0.0294 | 0.2344 ± 0.0117 | 0.1183 ± 0.0103 | 0.1449 ± 0.0173 | 0.1500 ± 0.0239 | 0.1550 ± 0.0100 | 0.0634 ± 0.0087 | 0.0873 ± 0.0127 | 0.0823 ± 0.0121 |
| CLEAN | `Unseen Test 10%` | 0.4689 ± 0.0135 | 0.3502 ± 0.0222 | 0.4073 ± 0.0299 | 0.3581 ± 0.0281 | 0.2468 ± 0.0122 | 0.1209 ± 0.0130 | 0.1704 ± 0.0180 | 0.1614 ± 0.0270 | 0.1813 ± 0.0102 | 0.0662 ± 0.0089 | 0.1033 ± 0.0123 | 0.0866 ± 0.0121 |
| EnzPlacer | `Unseen Test 10%` | 0.5270 ± 0.0139 | 0.3859 ± 0.0215 | 0.3782 ± 0.0199 | 0.4245 ± 0.0313 | 0.3034 ± 0.0130 | 0.1547 ± 0.0153 | 0.1775 ± 0.0198 | 0.1857 ± 0.0232 | 0.2260 ± 0.0112 | 0.0899 ± 0.0091 | 0.1130 ± 0.0111 | 0.1166 ± 0.0126 |

**Table S2. Detailed metrics for seen evaluations.** Mean ± half-width of the 95% bootstrap confidence interval for EC4 prediction performance (Accuracy, macro-F1, Precision, Recall) and hierarchical accuracies (EC1–EC3) on `Seen Test 50%`, `Seen Test 30%`, and `Seen Test 10%`.

| Model | Test set | EC4 Acc. | EC4 F1 | EC4 Prec. | EC4 Rec. | EC1 Acc. | EC2 Acc. | EC3 Acc. |
|---|---|---|---|---|---|---|---|---|
| BLAST | `Seen Test 50%` | 0.8971 ± 0.0071 | 0.7342 ± 0.0138 | 0.7541 ± 0.0137 | 0.7276 ± 0.0142 | 0.9651 ± 0.0047 | 0.9609 ± 0.0050 | 0.9571 ± 0.0052 |
| ProteInfer | `Seen Test 50%` | 0.7985 ± 0.0101 | 0.7270 ± 0.0124 | 0.7500 ± 0.0128 | 0.7210 ± 0.0122 | 0.8546 ± 0.0089 | 0.8378 ± 0.0087 | 0.8298 ± 0.0094 |
| GloEC | `Seen Test 50%` | 0.6557 ± 0.0127 | 0.4788 ± 0.0132 | 0.4914 ± 0.0141 | 0.5107 ± 0.0130 | 0.9270 ± 0.0067 | 0.8462 ± 0.0091 | 0.8028 ± 0.0103 |
| CLEAN | `Seen Test 50%` | 0.8775 ± 0.0084 | 0.7262 ± 0.0141 | 0.7363 ± 0.0141 | 0.7350 ± 0.0146 | 0.9772 ± 0.0039 | 0.9615 ± 0.0051 | 0.9474 ± 0.0056 |
| EnzPlacer | `Seen Test 50%` | 0.9098 ± 0.0074 | 0.7529 ± 0.0144 | 0.7601 ± 0.0147 | 0.7590 ± 0.0144 | 0.9827 ± 0.0033 | 0.9733 ± 0.0041 | 0.9658 ± 0.0046 |
| BLAST | `Seen Test 30%` | 0.5820 ± 0.0413 | 0.4185 ± 0.0384 | 0.4392 ± 0.0422 | 0.4101 ± 0.0393 | 0.6744 ± 0.0402 | 0.6660 ± 0.0412 | 0.6554 ± 0.0402 |
| ProteInfer | `Seen Test 30%` | 0.6561 ± 0.0369 | 0.5940 ± 0.0373 | 0.6150 ± 0.0373 | 0.5853 ± 0.0377 | 0.7247 ± 0.0369 | 0.7041 ± 0.0369 | 0.6921 ± 0.0369 |
| GloEC | `Seen Test 30%` | 0.3516 ± 0.0377 | 0.1933 ± 0.0274 | 0.2024 ± 0.0277 | 0.1967 ± 0.0280 | 0.8070 ± 0.0334 | 0.6492 ± 0.0386 | 0.6063 ± 0.0403 |
| CLEAN | `Seen Test 30%` | 0.6326 ± 0.0455 | 0.3765 ± 0.0390 | 0.3838 ± 0.0392 | 0.3815 ± 0.0384 | 0.9027 ± 0.0264 | 0.8397 ± 0.0339 | 0.7913 ± 0.0359 |
| EnzPlacer | `Seen Test 30%` | 0.6836 ± 0.0423 | 0.4016 ± 0.0394 | 0.4077 ± 0.0400 | 0.4073 ± 0.0400 | 0.9110 ± 0.0264 | 0.8587 ± 0.0328 | 0.8208 ± 0.0359 |
| BLAST | `Seen Test 10%` | 0.5608 ± 0.0665 | 0.4321 ± 0.0642 | 0.4594 ± 0.0659 | 0.4227 ± 0.0643 | 0.6305 ± 0.0616 | 0.6207 ± 0.0665 | 0.6158 ± 0.0665 |
| ProteInfer | `Seen Test 10%` | 0.6446 ± 0.0575 | 0.5933 ± 0.0552 | 0.6170 ± 0.0572 | 0.5826 ± 0.0547 | 0.7106 ± 0.0596 | 0.6915 ± 0.0617 | 0.6872 ± 0.0574 |
| GloEC | `Seen Test 10%` | 0.3851 ± 0.0618 | 0.2056 ± 0.0430 | 0.2125 ± 0.0439 | 0.2103 ± 0.0449 | 0.7873 ± 0.0554 | 0.6106 ± 0.0660 | 0.5851 ± 0.0617 |
| CLEAN | `Seen Test 10%` | 0.6371 ± 0.0616 | 0.3692 ± 0.0617 | 0.3809 ± 0.0625 | 0.3683 ± 0.0618 | 0.9068 ± 0.0395 | 0.8280 ± 0.0517 | 0.7692 ± 0.0591 |
| EnzPlacer | `Seen Test 10%` | 0.6757 ± 0.0640 | 0.3763 ± 0.0613 | 0.3866 ± 0.0630 | 0.3749 ± 0.0614 | 0.9061 ± 0.0419 | 0.8469 ± 0.0493 | 0.8073 ± 0.0542 |

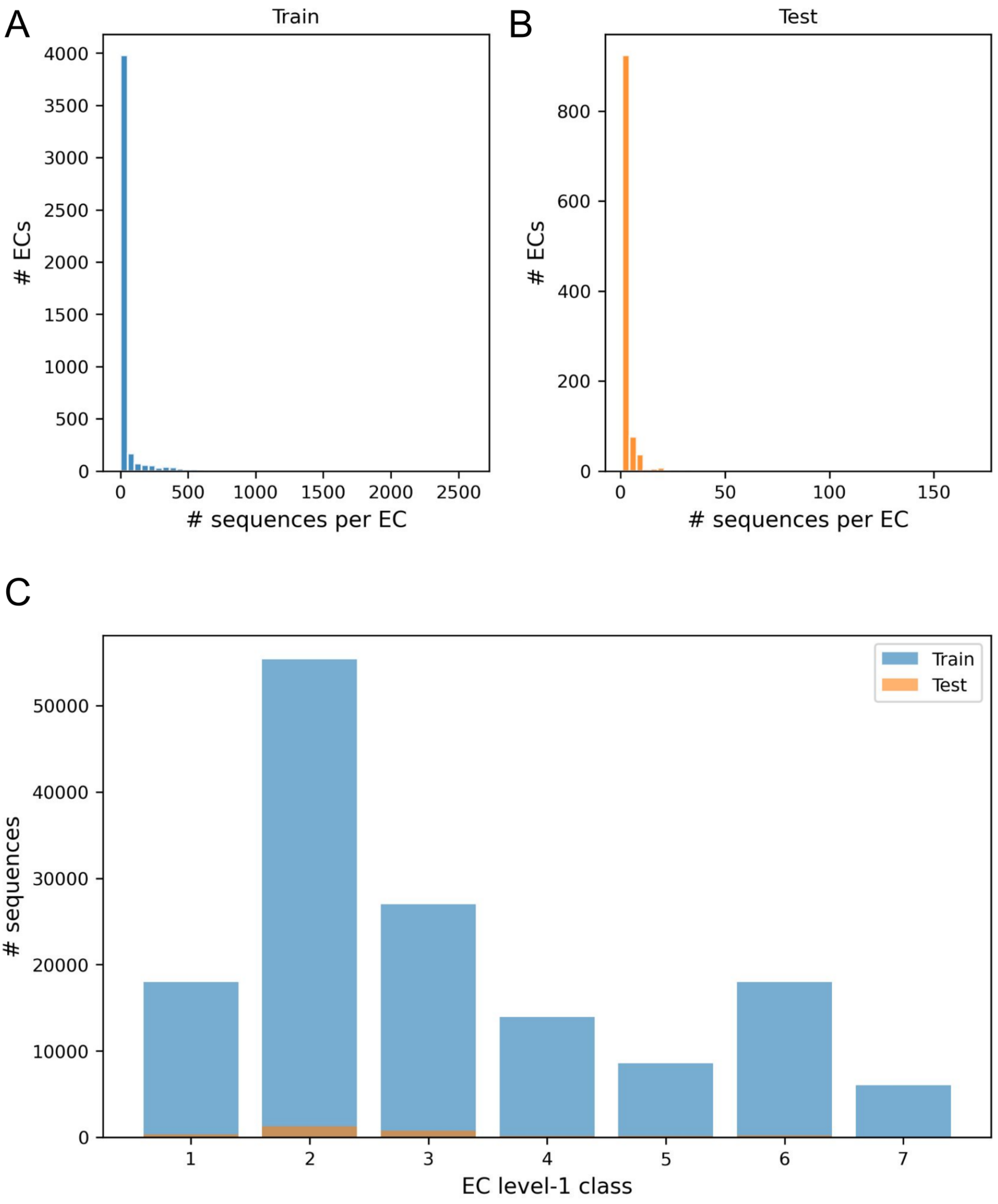


**Supplementary Figure S1. Seen dataset statistics.** (A–B) Distributions of the number of sequences per EC in the training and test splits. (C) Sequence counts per EC level-1 class for train and test.
